# The developing relations between networks of cortical myelin and neurophysiological connectivity

**DOI:** 10.1101/2020.07.07.191965

**Authors:** Marlee M. Vandewouw, Benjamin A.E. Hunt, Justine Ziolkowski, Margot J. Taylor

**Affiliations:** Department of Diagnostic Imaging, Hospital for Sick Children, Toronto, Canada; Program in Neurosciences and Mental Health, Hospital for Sick Children, Toronto; Department of Psychology, University of Toronto, Toronto, Canada; Department of Medical Imaging, University of Toronto, Toronto, Canada

**Keywords:** Magnetoencephalography, magnetic resonance imaging, connectivity, myelin, development

## Abstract

Recent work identified that patterns of distributed brain regions sharing similar myeloarchitecture are related to underlying functional connectivity, demonstrating cortical myelin’s plasticity to changes in functional demand. However, the changing relation between functional connectivity and structural architecture throughout child and adulthood is poorly understood. We show that structural covariance connectivity measured using T1-weighted/T2-weighted ratio and functional connectivity measured using magnetoencephalography exhibit nonlinear developmental changes. We then show significant relations between structural and functional connectivity, which have both shared and distinct characteristics dependent on the neural oscillatory frequency. Increases in structure-function coupling are visible during the protracted myelination observed throughout childhood and adolescence, and are followed by decreases near the onset of adulthood to potentially support increasing cognitive flexibility and functional specialization in adulthood. Our work lays the foundation for understanding the mechanisms by which myeloarchitecture supports brain function, enabling future investigations into how clinical populations may deviate from normative patterns.

## Introduction

One of the overarching goals of cognitive neuroscience is to identify biological markers of subtle mental and neurological illness, providing an objective means of assessment. Despite decades of research, very few clinically-useful discoveries have been made. One potential reason is that it is difficult to isolate atypicalities at the individual level without regular measurements during a period of typicality. A promising approach to alleviate this issue may be to isolate neural processes that are co-dependent and assess whether the dependency is altered during periods of pathology. In this regard, one process may act as the baseline, or period of typicality, for the other. Two obvious targets for such assessment are brain structure and brain function.

### Structure-function relations

One technique to assess the interplay between structure and function is to measure the brain’s structure (most commonly using magnetic resonance imaging (MRI)) and to take a proxy for function, such as behaviour. Examples include a series of studies that found associations between the behaviour of London taxi drivers and the properties of their hippocampi, including how long they had been navigating London’s streets (1) and those who passed a test requiring would-be taxi drivers to describe routes between different points in London (2). Another series of studies involved teaching participants new motor skills, such as juggling, and assessing changes in brain structure (3). These studies establish a directional link between behavioural change (a proxy for brain function) and brain structure, but do not provide a means of assessment in clinical populations.

### The interplay of structure and function

While the majority of studies assessing brain structure focus on grey matter properties such as cortical thickness, another readily quantifiable property is myelin. If the brain is compared to a bundle of wires, myelin, the sheath of oligodendrocytes encasing the axons of myelinated neurons, is the insulation. This insulation can increase the speed of transmission between neurons, and subsequently brain regions. However, myelin is not uniform across the brain, and complex processes result in selective myelination of neurons. Furthermore, the quantity of myelination differs between fibre tracts (4), suggesting a hierarchy of communication between distal regions.

In a seminal paper, Gibson and colleagues (5) found optogenetic stimulation of premotor cortex in a mouse model resulted in proliferation of oligodendrocyte precursor cells within the motor circuit. Moreover, examination of these same pathways four weeks later revealed increased quantities of new oligodendrocytes and thickness of the myelin sheath, as well as behavioural performance increases in the corresponding forelimb. This work provided the first direct demonstration of the mechanism by which brain function can shape brain structure, and underscores the plasticity of cortical myelin to changes in functional demand.

### Functional networks and structural connectivity

Modern cognitive neuroscience rarely considers individual brain regions in isolation, but recognises that most brain structures operate as part of functional networks. These networks are now generally accepted to underpin brain function in the widest sense, from motor movement to higher-order cognitive processes. Magnetoencephalography (MEG) is especially well suited for direct assessment of these functionally coupled networks. MEG measures the minute magnetic fields originating from the dendrites of communicating populations of pyramidal neurons. Measurement of these fields, detected noninvasively outside of the skull, combined with appropriate mathematical modelling, result in a technique with good spatial resolution and very high temporal resolution. The oscillating fields detected by MEG occur at a range of frequencies, and functional networks can exist within a single frequency band or across multiple spectral bins. Different frequency bands are associated with distinct processes, such as beta band with movement and alpha with memory (6, 7).

As we understand brain function to operate on a network level, so too would we anticipate that brain structure covaries across regions, forming networks. The primary means to evaluate this is by assessment of structural covariance over subjects (8, 9). Regions that form part of the same structural network would be expected to have a high degree of similarity in the ratio of a structural property over subjects. For example, a common finding is that structural properties within Wernicke’s Area correlate, across subjects, with the same structural properties in Broca’s Area, forming hubs of a language network (10).

In children, evidence indicates that some functional networks are present before birth (11), while others continue to mature well into adulthood (12, 13). Furthermore, the findings of the taxi driver and juggling studies would suggest that as an individual becomes more uniquely specialised, the brain’s structure will evolve to support that specialised behaviour; the brain structure of a juggling taxi driver would likely be distinct in motor and hippocampal regions from either taxi drivers or jugglers only. In turn, this specialised structural connectivity underpins unique functional networks (14) that would be similarly distinct between individuals. These two discrete connectomes would result in a more proximal function-structure relationship compared to less specialised individuals. As the amount of behavioural and cognitive specialisation generally increases with age, we would hypothesise that the structure-function relations also increase with age.

### The relation between functional connectivity and structural covariance of cortical myelin

Previously (15), we investigated the relation between cortical myelin and functional connectivity, as measured by MEG, in healthy adults (N=58; aged 18-65 years). We found a relation between estimates of cortical myelin and MEG-derived functional connectivity measurements that varied on a spectral basis. We observed that beta (∼13-30Hz) and low-gamma (∼30-70Hz) frequency functional connectivity matrices more closely related with the structural covariance of cortical myelin. However, due to the size of our sample, we were unable to assess how the proximity of this relationship changes with age.

Furthermore, cortical myelin was measured using Magnetization Transfer (MT) sequences (16) within a 7-Tesla MRI. While these sequences are robust, they are highly specialised and often not feasible at lower field strengths, and acquisition times are lengthy. An alternative approach is to take the ratio of two commonly acquired images, T1-weighted (T1-w) and T2-weighted (T2-w) to form a voxel-wise map of myelin estimates (17–20). The concordance between MT and the T1-w/T2-w ratio appears to be high in both the white and grey matter (21), and has been shown as a valid and reproducible measure to use when examining structural covariance (22). Moreover, as the two elements required for this technique are often routinely acquired, there are opportunities for retrospective analyses of large datasets. We therefore assessed function-structure relations in a large sample spanning young children through to adults using T1-w/T2-w ratio estimates of cortical myelin to establish for the first time foundational developmental data on this relationship.

### Aims and Hypotheses

We investigated how the relation between brain function and structure changes as humans age. We hypothesised the relation between structure and function to increase in strength with age. We anticipated this increase to be most apparent within the beta and low gamma band, where we previously saw the greatest relation between structure and function in adults.

## Results

### Marked connectomic differences between the youngest and oldest age groups

One-hundred and ninety-six participants aged between four and 45 years (M = 19.95 years, SD = 11.16 years, 130 males) are included in the current study, who were recruited as part of several projects in the Taylor lab at the Hospital for Sick Children (Toronto, Canada). Myelin was measured by calculating a T1-w/T2-w ratio image for each participant. For each of the 90 regions in the Automated Anatomical Labelling (AAL) atlas (23), myelin intensity was calculated as the median intensity along the surface midway between the grey and white matter. Five minutes of resting-state MEG data were also obtained from each participant. For each of the canonical frequency bands (theta (θ), alpha (α), beta (β), low gamma (γ_1_), and high gamma (γ_2_)), amplitude envelope connectivity (AEC) was calculated between pairwise AAL regions, measuring functional connectivity. A sliding window approach was employed to compute the structural covariance and mean AEC for each frequency band for age windows spanning 6.9 – 32.7 years. The structural covariance and mean AEC networks are shown for the youngest and oldest age windows in Figure 1A and Figure 1B-F, respectively.

**Figure 1.**
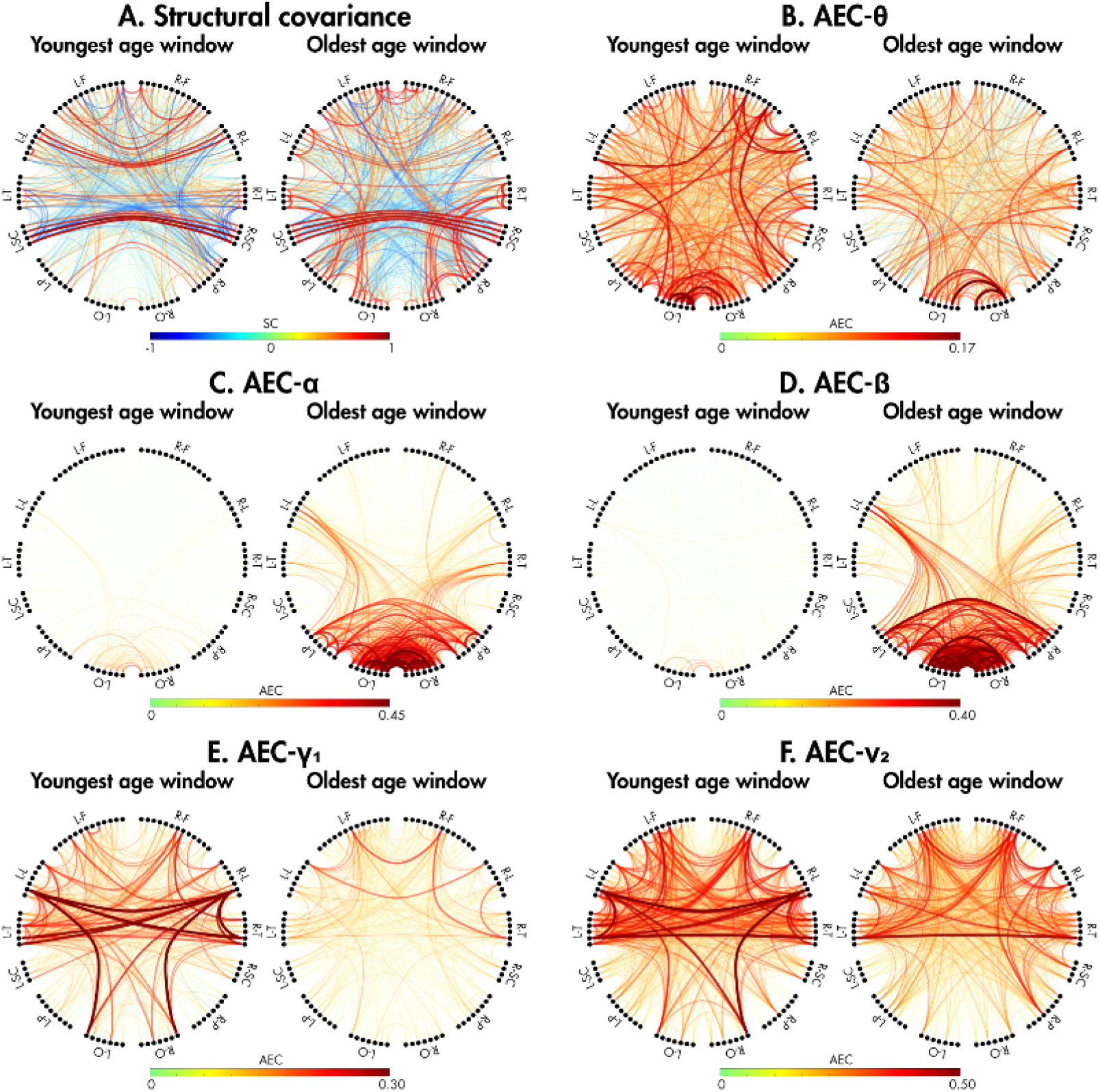
Structural covariance (A) and functional connectivity, measured using AEC (θ: (B); α: (C); β: (D); γ_1_: (E), and γ_2_: (F)), are shown for the youngest (median age: 6.89 years) and oldest (median age: 32.68 years) age windows.

For each connection in the structural and functional network, linear and nonlinear (locally adaptive smoothing splines) fits were applied to examine the relation between connectivity (structural covariance or AEC) and the median age at each time window. Connections were classified as linear (yellow) or nonlinear (purple) based on the fit with the minimum Akaike’s Information Criteria (AIC), excluding non-significant connections (white, *p*_corr_≥0.05). In all cases, at least 80% of the connections were classified as having a nonlinear relation with age, with 1-2% classified as having a linear relation (Table 1). For both the structural covariance (SC) and functional networks, the effective degrees of freedom for the nonlinear spline fits was always approximately 3.5, meaning trajectories were quadratic.

**Table 1.** Percent of connections classified as having a linear, nonlinear, or non-significant relation with age.

|  | % Connections |  |  |
| --- | --- | --- | --- |
|  | Linear | Nonlinear | Not significant |
| **SC** | 1.1 | 85.6 | 13.4 |
| **AEC-θ** | 1.4 | 88.7 | 10.0 |
| **AEC-α** | 1.5 | 89.0 | 9.5 |
| **AEC-β** | 1.6 | 88.9 | 9.5 |
| **AEC-$\gamma_1$** | 2.0 | 85.4 | 12.6 |
| **AEC-$\gamma_2$** | 2.0 | 85.7 | 12.3 |

### Structural and functional connectivity peaks in different age windows

After fits were classified as linear or nonlinear, connectivity was fitted using the classified fits at 200 equally spaced ages spanning the age range (between 6.9 and 32.7 years, at intervals of 0.19 years). The age at which the maximum fitted SC or AEC was reached (Age(SC_max_) or Age(AEC_max_)) was recorded (Figure 2A and Figure 2B-F, respectively). The darkest red connections indicate that maximum connectivity occurred near the last age window and thus exhibited an overall positive relation with age, while the darkest blue connections indicate that maximum connectivity occurred in the first age window, exhibiting an overall negative relation with age. A breakdown of the percentage of significant connections for each connectivity network with Age(SC_max_) or Age(AEC_max_) falling in different age brackets is presented in Table 2. For structural covariance (Figure 2A), the majority of connections have an Age(SC_max_) occurring near the first or last age window (<8 years or >30 years). Connections between left parietal and occipital regions to right limbic, temporal, and subcortical regions decreased over our age span; connections among the subcortical structures increased with age, while the remaining connections tended to decrease. In contrast, the Age(AEC_max_) for the five canonical frequency bands were more dispersed across the age span. The pattern for AEC in the theta band (Figure 2B) included connections within the parietal lobes peaking around 20 years of age, and within the right limbic structures peaking early in childhood. In alpha and beta (Figure 2C and 2D, respectively), approximately 45% and 55%, respectively, of connections reached their peak near the end of our age range (>30 years), with the majority of connections to the frontal lobe, particularly in the right hemisphere, reaching maximum connectivity earlier in life. For both gamma bands (Figure 2E and 2F), about 40% of the connections reached their maximum near the first time window (<8 years), implying that connectivity decreases with age, with the exception of connections extending from the bilateral parietal lobes.

**Table 2.** Percentage of connections with the age of maximum fitted connectivity (Age(SC_max_/AEC_max_)) occurring in different age brackets.

| | % Significant connections with $\text{Age}(\text{SC}_{\max}/\text{AEC}_{\max})$ | | | | |
| --- | --- | --- | --- | --- | --- |
|  | < 8 years | 8 – 12 years | 13 – 19 years | 20 – 30 years | > 30 years |
| **SC** | 37.9 | 2.4 | 9.1 | 13.4 | 37.3 |
| **AEC-θ** | 33.3 | 22.9 | 26.7 | 23.8 | 24.0 |
| **AEC-α** | 18.9 | 28.4 | 33.8 | 35.7 | 45.4 |
| **AEC-β** | 24.0 | 26.4 | 31.2 | 37.0 | 56.7 |
| **AEC-$\gamma_1$** | 42.9 | 20.2 | 23.6 | 25.4 | 34.7 |
| **AEC-$\gamma_2$** | 42.9 | 20.1 | 22.8 | 23.6 | 26.3 |

**Figure 2.**
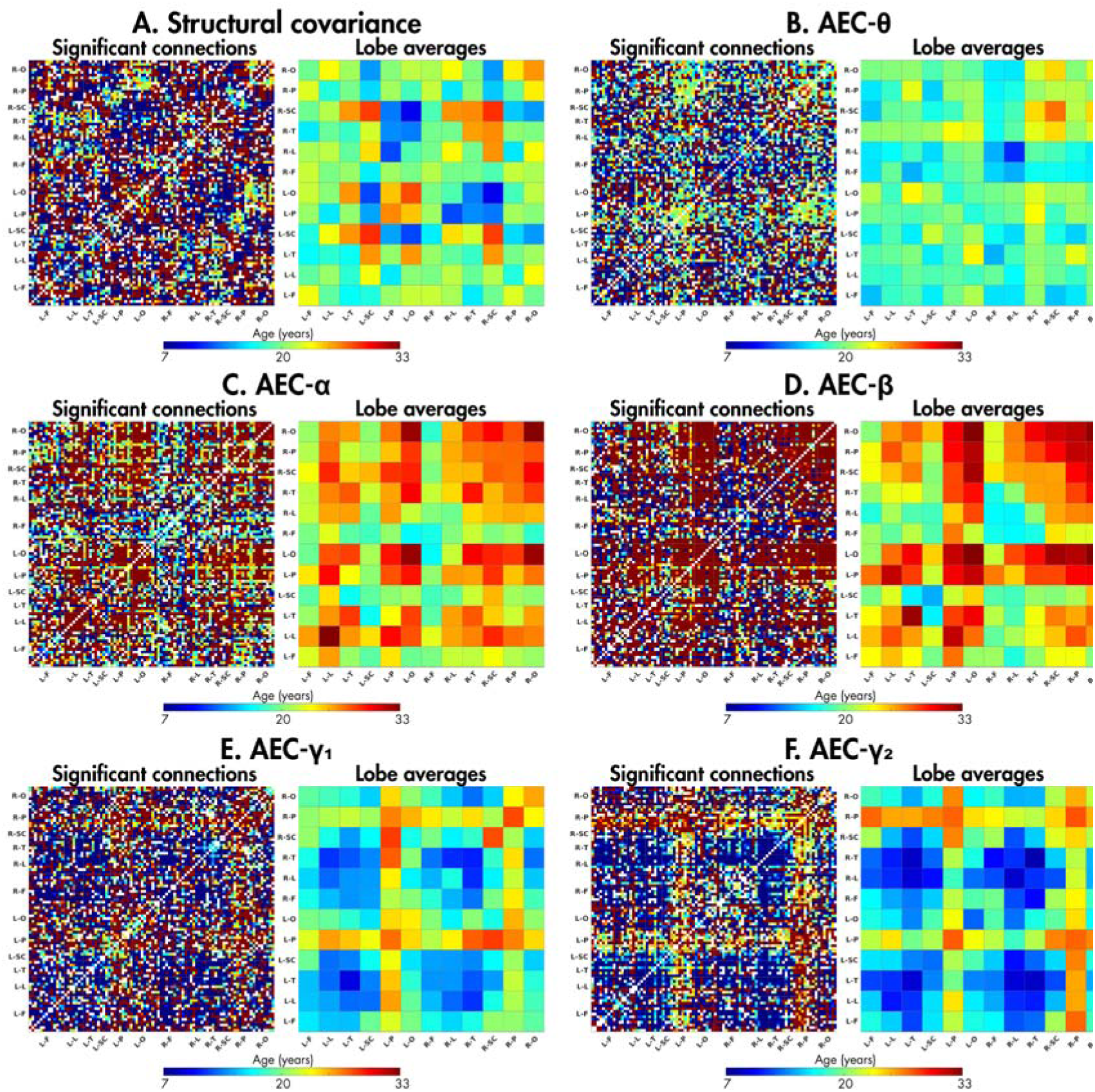
For connections with significant age-related trajectories, the ages at which the fitted structural covariance (Age(SC_max_); A) and AEC (Age(AEC_max_); θ: (B); α: (C); β: (D); γ_1_: (E), and γ_2_: (F)) reached a maximum are shown; non-significant connections are indicated by a white matrix element). Data averaged over connections within and between lobes are also presented. The darkest red connections indicate that the fitted connectivity reached a maximum near the last age window and thus showed an overall positive relation with age, while the darkest blue connections indicate that the fitted connectivity reached a maximum near the first age window and thus exhibited an overall negative relation with age.

### The magnitude of connectivity change varies significantly across the cortex

The maximum change in fitted structural covariance (ΔSC) and AEC (ΔAEC) along the trajectory was also calculated (Figure 3A and Figure 3B-F, respectively). Similar to Figure 2, blue connections indicate an overall decrease in connectivity across our age span, while red connections indicate an overall increase. In all cases, regions with fitted connectivity reaching their maximum early or late in the age span demonstrated the greatest change in connectivity across the age span, seen by comparing the lobe averages in Figures 2 and 3. The percentage of all significant connections with positive and negative trajectories (indicating increased connectivity or decreased connectivity with age, respectively) are shown in Table 3; the percentages are also shown when only considering connections whose absolute change in connectivity is in the top 10% of all significant connections.

**Table 3.**
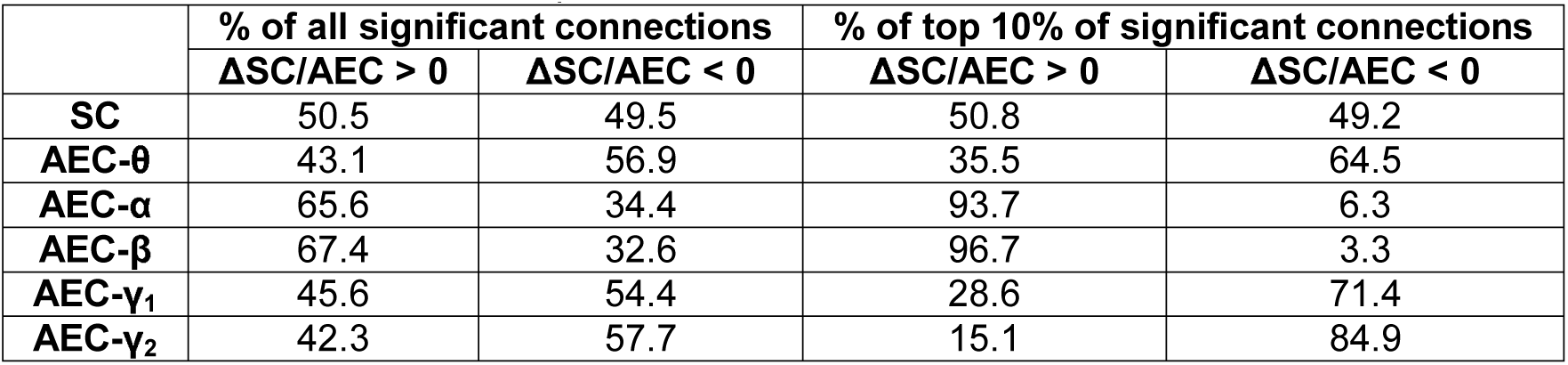
Percent of connections with positive and negative maximum change in connectivity (ΔSC or ΔAEC), indicating overall increased or decreased connectivity with age, respectively. Data are first shown for all connections, followed by only considering the connections whose absolute ΔSC/AEC falls in the top 10% of all connections.

**Figure 3.**
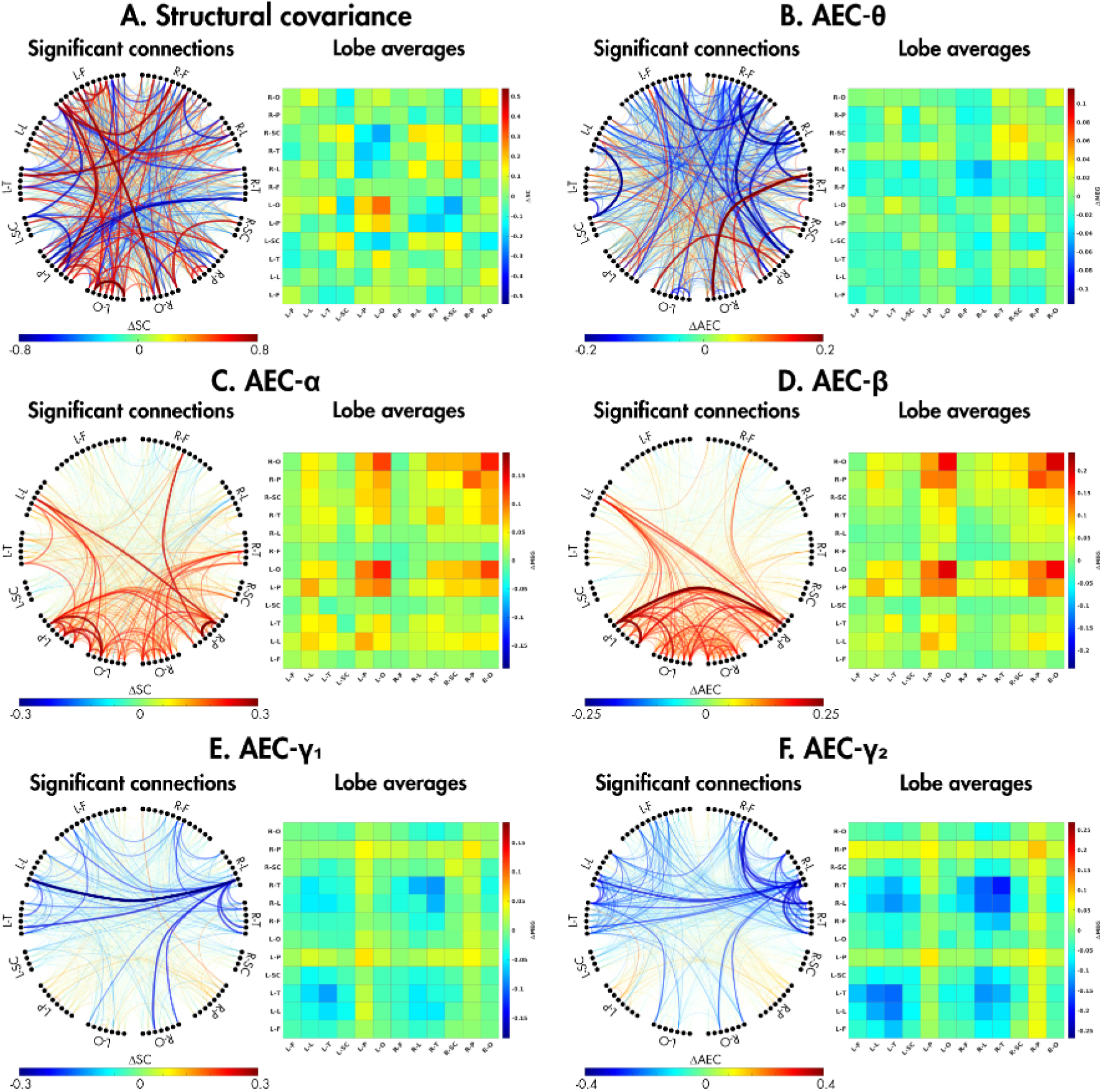
For connections with significant age-related trajectories, the maximum changes in structural covariance (ΔSC; A) and AEC (ΔAEC θ: (B); α: (C); β: (D); γ_1_: (E), and γ_2_: (F)) are shown; data averaged over connections within and between lobes are also presented.

### The developmental relation between structure and function is spectrally distinct

For structural covariance, there appears to be no trend in the directionality of the maximal change in connectivity, with about 50% of all connections being positive, even when only the top 10% of connections are considered. A similar trend is found for AEC in the theta band; however, when considering only the connections whose ΔAEC is in the top 10%, the change in connectivity over age shifts to be more positive. In both alpha and beta, about two thirds of all connections display a positive change in AEC across age, with this proportion shifting to about 95% when only considering the top connections. Conversely, in low and high gamma, the majority of connections show a negative change in AEC across age, with greater than 70% of the top connections showing this pattern.

Finally, and most importantly, fitted structural covariance was correlated across all connections with the functional connectivity in each frequency band to examine changes with age (Figure 4). In all bands, the structure-function correlation was significant across the age range. When examining the changing relation between structure and function in the theta band, the structure-function correlation increased during childhood, peaking at 15 years of age before beginning to decline. Structure-function coupling was less strong in alpha and beta compared to theta, with the structure-function relation strength increasing in childhood, peaking at 16 and 15 years, respectively, before remaining relatively stable with a slight decline. Low gamma showed the most dramatic change in structure-function correlation across age: correlation strength approximately doubled between 7 and 22 years of age, with its peak in young adulthood exhibiting the strongest structure-function coupling across all frequency bands. Finally, in high gamma, the structure-function coupling became stronger during childhood, reaching its maximum strength at 17 years before remaining relatively stable but with a slight decline, which accelerated at around 27 years of age.

**Figure 4.**
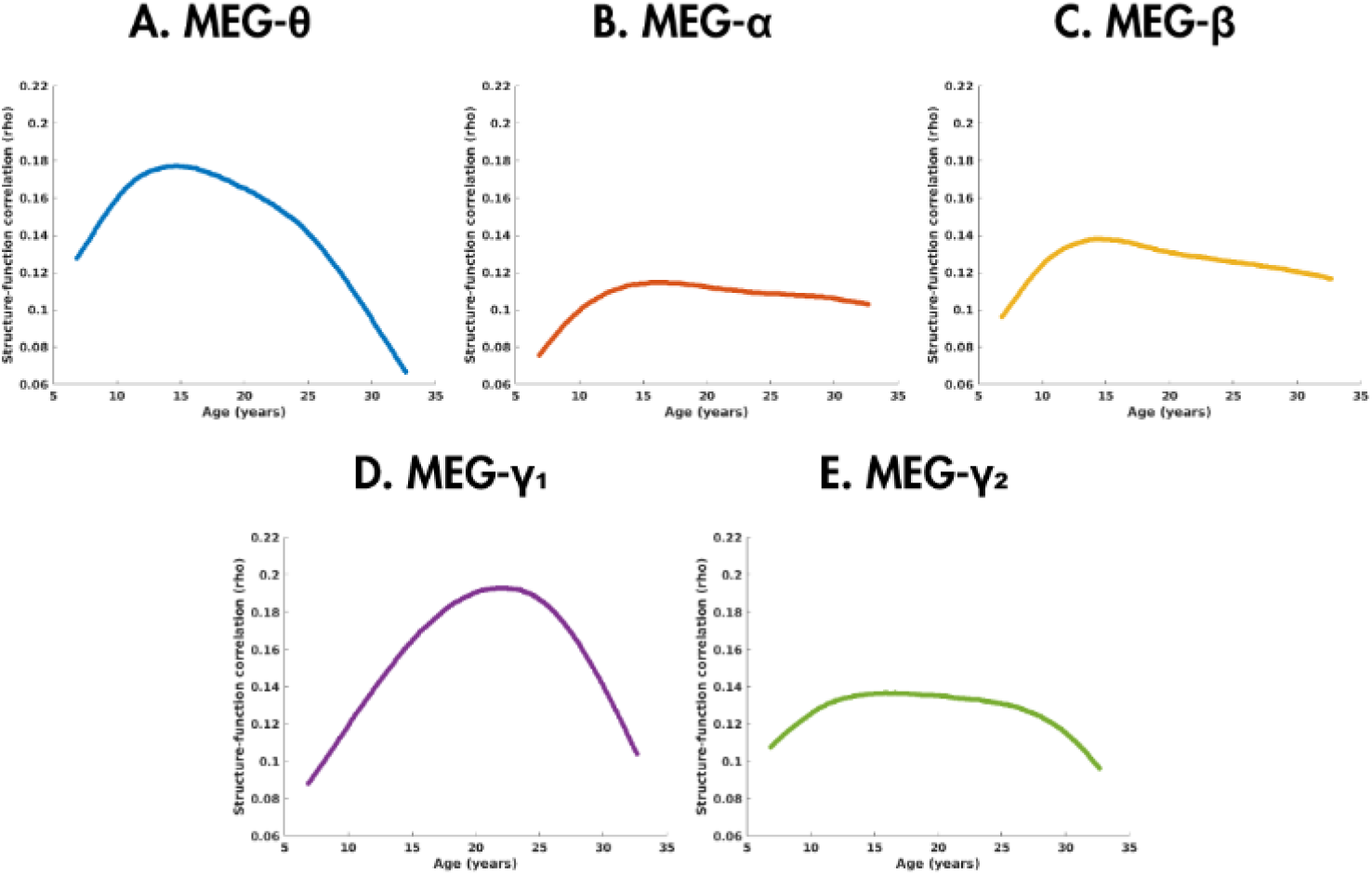
Age-related changes in the correlation (Spearman’s rho) between structural covariance and functional connectivity (AEC) in the five canonical frequency bands (θ: A; α: C; β: D; γ_1_: E, and γ_2_: F).

## Discussion

Quantifying the relation between brain structure and function offers a unique avenue of exploration to identify putative biomarkers of psychopathology. As an initial step, we first assessed the viability of T1-w/T2-w ratio for use in the estimation of structural connectivity. Second, we investigated how both structural covariance and functional connectivity, as measured by T1-w/T2-w and AEC, respectively, changed with age. Most importantly, we assessed how the relation between structural and functional networks changed with age.

Typically, cortical myelin is measured using MT sequences as in our previous papers (15, 16); however, the necessary sequences are complex and best used on a 7T MRI. T1-w and T2-w images are routinely collected as part of image acquisition protocols, and their ratio has been established as a good approximation of myelin (17), opening the door to retrospective studies. The use of T1-w/T2-w ratio in measuring structural covariance has been shown to produce brain organization similar to that of other structural measures and is validated as reproducible in adults and children (21, 22). Our data support the use of T1-w/T2-w ratio to measure myelin covariance – our structural covariance networks demonstrated organization as well as high and symmetric intra-hemispheric connectivity in both children and adults, consistent with our previous study using MT (15). Myelin covariance is highest between the bilateral subcortical and limbic structures, both within and between hemispheres, and amongst within-hemispheric connections in the occipital and parietal lobes (particularly in the adults), as seen by Melie-Garcia et al. (24).

Consistent with previous studies examining typical development from childhood to young adulthood, the age-related changes of both structural covariance (24–27) and MEG functional connectivity (28–30) follow predominantly nonlinear, quadratic trajectories. The differences in functional connectivity with age was marked by increases in functional connectivity with age in the alpha and beta bands and decreases in the gamma bands, consistent with developmental changes reported in both phase-based (31) and amplitude (30, 32) connectivity measures. Although individual connections were found to change significantly with age in the theta frequency band, the changes exhibited no clear pattern or network organization, aligning with previous findings (29, 31).

In each frequency band, however, there was a relation between structural covariance and functional connectivity that remained significant across the age span. In all bands, the function-structure relation strength increased with age, before beginning either a slow or rapid decline, contrary to our hypothesis of consistently increasing relation strength. Humans experience protracted myelination that extends into adulthood (33). The observed increases in structure-function coupling during adolescence could reflect the ongoing reorganization of underlying myeloarchitectural networks to support functional activity, or, due to the linked nature between myeloarchitecture and functional connectivity, structural reorganization could shape functional connectivity. Once the window of prolonged myelin maturation has passed, the observed decreases in structure-function coupling could be supporting cognitive flexibility and functional specialization, as past studies have demonstrated that on a regional level, structure-function coupling is inversely associated with the brain region’s functional complexity (34–36). Furthermore, as brain regions develop distinct specialisation, the across-brain variance in the structure-function relation strength would increase, and thus decrease whole-brain structure-function coupling.

Low gamma exhibited the strongest relationship between structure and function, as hypothesized, with marked increases in correlation strength during childhood and adolescence, before demonstrating a decrease in relation strength at the onset of adulthood. It has been proposed that high frequency oscillations such as gamma are generated by local neuronal networks (37, 38); thus, underlying structure is vital for the long-range integration of distributed brain regions in the gamma frequency band. Furthermore, theta oscillations are believed to integrate local gamma oscillations into large-scale brain networks (37), and theta band activity can modulate gamma band activity during cognition (39, 40). The interplay between theta and gamma band oscillations could explain the similar age-related changes in structure-function coupling in these two bands, with the low gamma band lagging behind theta, which could be due to local detailed processing continuing to mature after the long-range connections are established.

Gamma activity is evoked or induced by a wide range of sensory and cognitive processes and gamma synchronisation is considered of fundamental importance in processes throughout the brain (41), dependent upon excitatory/inhibitory balance (42). Gamma has been frequently associated with local processing or communication as well as being critical for inter-layer or inter-regional communication, organising neural activity in combination with lower frequencies. However, there is still uncertainty regarding the origins of gamma and the distinct roles for low and high gamma (which are variously defined) (for reviews, see (43, 44)). Contributing to this ongoing field of study, our findings that high and low gamma connectivity showed similar age-related changes, yet differences in their relation to structural covariance was marked, underscores the importance of investigating these frequency bands independently.

Although we also hypothesized that the beta band would show a strong structure-function relation, its peak at around 15 years of age was low relative to the peaks in the theta and low gamma bands, showing a similar flattened age-related trajectory to alpha. The mean age in our original work from which our hypothesis was derived was 39 years old (15), however, and therefore beyond our current age span; yet, at our oldest age window (33 years of age), beta did, in fact, demonstrate the highest structure-function correlation strength relative to the other four frequency bands. Given the age-related trajectories of the bands, we would predict that the structure-function coupling in beta would remain high relative to the other bands in older adulthood, coinciding with our initial hypothesis. Beta plays a strong role in coordination among sensory and motor inputs and is particularly important for cognitive top-down control (44). Although many other roles have been suggested for beta, even these suggest that it is a critical brain rhythm that would remain strong and stable over adulthood.

Alpha, after the initial rapid development increase, showed the most stable structure-function relation, closely followed by beta. Both frequency bands also had similar posterior-dominated functional connectivity patterns and age-related trajectories. Both frequencies are implicated in long-range brain connectivity with greater involvement of deep cortical layers for beta (45, 46). As alpha has been linked with a range of critical cognitive functions, such as memory, inhibition and cognitive control (47–49), this stability may reflect the ongoing utilisation of these functions through to mid-adulthood.

Having established how the relation between brain function and structure changes as humans age, a natural future direction is to investigate whether clinical samples show similar patterns in structure-function coupling, or whether they exhibit deviated age-related trajectories. Disorders exhibiting connectivity atypicalities in both brain structure and function, such as autism spectrum disorder, would be of particular interest. Furthermore, expanding the age range in the current study would give insight into whether structure-function relations stabilize in mid-adulthood and allow for the investigation into whether or not these relations deteriorate during aging.

## Materials and Methods

### Participants

Data collected on several projects within the Taylor lab at the Hospital for Sick Children (Toronto, Canada) were pooled into a single database of 425 individuals based upon the existence of resting state MEG. To be included in the current analysis, participants were required to be free of contraindications for MEG or MRI, and a history of/current neurological disorders and/or mental illness. In addition, all participants were required to have completed a 5-minute resting state acquisition with static fixation cross in MEG, and useable T1- and T2-weighted images obtained using a 3T MRI. Following exclusion based on the aforementioned criteria, a cohort of 207 individuals was compiled. Following closer inspection of MRI data during FreeSurfer segmentation and upon calculation of T1-w/T2-w ratio, a further 11 participants were removed due to excessive movement in the scanner preventing accurate segmentation (n = 5) or T1-w/T2-w ratio values in 2 or more regions falling >3 standard deviations from the group mean (n = 6). The final cohort consisted of 196 participants (M: 19.95 years, SD = 11.17 years, 130 males).

### Data acquisition

Five minutes of resting-state MEG data were acquired in supine position using a CTF 151-channel system (CTF-MISL, Coquitlam, Canada) positioned within a magnetically shielded room. A centrally positioned grey cross within a circle on a black background was back projected onto a screen inside the MEG suite, and participants were instructed to fixate on the cross. Data were acquired at a sampling frequency of 600 Hz operating in third-order synthetic gradiometry configuration. Participants were fitted with three head position indicator coils, located at the nasion and left and right preauricular points, to measure moment-to-moment head motion. The location of these coils was recorded, and MRI-visible markers were positioned at these locations to enable coregistration between MEG and MRI data

MRI data were acquired on a 3.0T Siemens Trio (12 channel head coil) or a PrismaFIT (20 channel head and neck coil). T1-w images were collected using 3D magnetization prepared rapid gradient echo (MPRAGE) sequences (Trio: TR/TE=2300/2.96ms, FA=9°, FOV=240×256mm, number of slices=192, resolution=1.0mm isotropic, scan time=5:03min; PrismaFIT: TR/TE=1870/3.14ms, FA=9°, FOV=240×256mm, number of slices=192, resolution=0.8mm isotropic, scan time=5:01min). T2-w images were collected using 2D turbo-spin echo (TSE) sequences (Trio: TR/TE=9000/104ms, FA=120°, FOV=191×230mm, number of slices=128, resolution=1.2mm isotropic, scan time=2:42min; PrismaFIT: TR/TE=3200/408ms, FA=120°, FOV=256×256mm, number of slices=192, resolution=0.8mm isotropic, scan time=6:11min).

### MRI processing and intra-cortical myelin mapping

Participants’ T1-w and T2-w images were used for cortical reconstruction and segmentation using the FreeSurfer image analysis suite, which is documented and freely available for download (http://surfer.nmr.mgh.harvard.edu/, version 6.0), and whose technical details are described in previous publications (50–61). Manual editing was performed for every scan to ensure accuracy of the pial surface and grey-white matter boundary. Once quality standards were met, a mask of the mid-surface, halfway between the grey and white matter, was extracted.

To calculate the T1-w/T2-w ratio image for each participant, the procedure outlined in (62) was employed. The T2-w image was linearly registered to the T1-w image using FMRIB’s linear image registration tool (63, 64), and both images were corrected for bias field inhomogeneities (65). The image intensities were calibrated using non-tissue masks in the eye and temporal muscle. The calibrated T1-w image was then divided by the calibrated T2-w image to obtain a T1-w/T2-w ratio image for each participant.

Each participant’s T1-weighted image was nonlinearly co-registered to MNI152 standard space (66, 67) and the generated transformations were used to register the AAL atlas into participant space. The AAL atlas was then masked with the mid-surface and applied to the T1-w/T2-w ratio image, and the median T1-w/T2-w ratio value of each AAL parcel along the mid-surface was extracted. This generated an array of 90 values per participant, representing the varying intensity of cortical myelin across the brain.

### MEG processing and Amplitude Envelope Connectivity

MEG data were bandpass filtered between 1-150Hz, and notch filters were applied at 60 and 120Hz. An independent component analysis (ICA) was then performed to remove artefacts arising from the heart and eyes (68). Next, an automated data rejection algorithm was used to detect segments of data containing excessive head movement (>7 mm) or artefact. The remaining data were epoched into as many 10-s “trials” as available. Note that these trials are arbitrary divisions of a continuous resting-state recording and are epoched for computational reasons. Participants with greater than five trials remaining following this procedure were taken forward for primary analyses (31).

A linearly constrained minimum variance (69) spatial filter, implemented in Field Trip (70), was used to enter source space. The forward model was based upon a dipole approximation (71) and a realistically shaped single-shell approximation (72). Dipole orientation was determined using the maximum eigenvector and fixed across trials. Covariance was calculated within a 1– 150Hz covariance window spanning the entire experiment and regularized using the Tikhonov method (73) with a regularization parameter equal to 5% of the maximum eigenvalue of the unregularized matrix. The beamformer output was normalized by estimated noise, the result being termed the neural activity index, to avoid biasing measurements toward the centre of the head. Participant data were transformed onto the 90-parcel Automated Anatomical Labelling (AAL) atlas (23) and a beamformer spatial filter was used to derive a single signal from each region at the region’s centre of mass.

Amplitude envelope connectivity (AEC) was calculated by first filtering continuous source-space data into canonical frequency bands (theta (θ; 4-7Hz), alpha (α; 8-14Hz), beta (β, 15-30Hz), low gamma (γ_1_; 31-55Hz), and high-gamma (γ_2_; 65-120Hz)) and subsequently reducing source leakage by orthogonalizing data using the Colclough technique (74), implemented within the “MEG-ROI-Nets” toolbox (https://github.com/OHBA-analysis/MEG-ROI-nets). The absolute of the Hilbert transform (or the amplitude envelope) was then calculated on a source-wise basis. Envelope data were demeaned and down-sampled to 1Hz by averaging over samples within the 10s epochs. Subsequently, whole brain AEC was determined by calculating, for each participant and frequency band, the Pearson correlation coefficient for each regional interaction, generating a 90-by-90 symmetric adjacency matrix.

### Network construction

To construct age-dependent networks, a sliding window approach was employed (24, 25). The number of participants per window (window width) was held constant at 50, well above the minimum number of participants necessary for reliable correlations when calculating structural covariance (75). The first age window consisted of the 50 youngest participants, and the youngest two participants were replaced with the next two oldest participants in each subsequent window (meaning a step size of two). The “age” for each window was defined as the median age for all participants included, and ranged from 6.89 – 32.68 years. To ensure results were not influenced by the window width or step size, all subsequent analyses were repeated with window widths of 40 and 60 and step sizes of one and five (see Supplemental Information for further details).

To derive a network of cortical myelin intensity, we calculated structural covariance by computing the correlation (Spearman, due to non-normality), of T1-w/T2-w ratio values across window-specific participants, generating a 90-by-90 symmetric adjacency matrix for each age window. Prior to correlating, a linear regression was performed on the raw T1-w/T2-w ratio values to remove the within-window effects of sex and acquisition scanner (PrismaFT/Trio). For each frequency band, MEG functional connectivity networks for each age window were computed by averaging the AEC adjacency matrices across the window-specific participants after removing the within-window effect of sex.

### Assessing age-related changes

For every unique connection (4,005 connections: 90-by-90 symmetric matrix, excluding diagonal), the age-related trajectory of structural covariance was fitted using a linear and locally adaptive smoothing spline (nonlinear; (25, 26, 76)) fits. The smoothing spline was computed as a weighted sum of six *B*-spline basis functions constrained to be at least as smooth as a cubic fit with knots equally spaced across the median age of the windows and optimal smoothing determined using restricted maximum likelihood (76). For both the linear and nonlinear fits, False Discover Rate (FDR; (77)) correction was performed on the resulting *p*-values to control for multiple comparisons across each connection, and significance was held at *p*_corr_<0.05. Linear and nonlinear fit quality was evaluated using Akaike’s Information Criteria (AIC); connections with a lower linear AIC than nonlinear AIC were classified as linear (if significant), and vice versa. After classification, structural covariance trajectories were fitted at 200 equally spaced ages between the minimum (6.89) and maximum (32.68) median age. For significant linear and nonlinear fits, the age at which the fitted structural covariance was at its maximum (Age(SC_max_)) was recorded, along with the maximum change in structural covariance (ΔSC) along the trajectory. For each frequency band, age-related trajectories of AEC were determined using an identical procedure.

For each frequency band, the relation between function and structure was assessed by correlating (Spearman’s, due to non-normality) the fitted structural covariance and functional connectivity across all connections at each of the 200 fitted points. Resulting *p*-values were FDR corrected across the 200 correlations and significance was held at *p*_corr_<0.05. Spearman’s rho values were plotted against age to examine the changes in strength of the structure-function relations.

## Acknowledgments

We thank the following individuals who were involved in the data acquisition for this study: Rachel Leung, Vanessa Vogan, Sarah Mossad, Julie Sato, Veronica Yuk, MyLoi Huynh, and Julie Lu, as well as our wonderful MRI and MEG technologist: Ruth Weiss, Tammy Rayner and Marc Lalancette. We also would like to thank the participants and their families for their invaluable contribution.

## References

1. E. A. Maguire, et al., Navigation-related structural change in the hippocampi of taxi drivers. Proc. Natl. Acad. Sci. U. S. A. 97, 4398–4403 (2000).

2. K. Woollett, E. A. Maguire, Acquiring “the Knowledge” of London’s Layout Drives Structural Brain Changes. Curr. Biol. 21, 2109–2114 (2011).

3. J. Scholz, M. C. Klein, T. E. J. Behrens, H. Johansen-Berg, Training induces changes in white matter architecture. Nat. Neurosci. 12, 1370–1371 (2009).

4. C. Lebel, et al., Diffusion tensor imaging of white matter tract evolution over the lifespan. Neuroimage 60, 340–352 (2012).

5. E. M. Gibson, et al., Neuronal Activity Promotes Oligodendrogenesis and Adaptive Myelination in the Mammalian Brain. Science. 344, 1252304 (2014).

6. A. Fry, et al., Modulation of post-movement beta rebound by contraction force and rate of force development. Hum. Brain Mapp. 37, 2493–2511 (2016).

7. J. M. Palva, S. Monto, S. Kulashekhar, S. Palva, Neuronal synchrony reveals working memory networks and predicts individual memory capacity. Proc. Natl. Acad. Sci. U. S. A. 107, 7580–7585 (2010).

8. O. Nempont, J. Atif, E. Angelini, I. Bloch, Combining Radiometric and Spatial Structural Information in a New Metric for Minimal Surface Segmentation BT - Information Processing in Medical Imaging in N. Karssemeijer, B. Lelieveldt, Eds. (Springer Berlin Heidelberg, 2007), pp. 283–295.

9. O. Coulon, et al., Structural group analysis of functional activation maps. Neuroimage 11, 767–782 (2000).

10. J. P. Lerch, et al., Mapping anatomical correlations across cerebral cortex (MACACC) using cortical thickness from MRI. Neuroimage 31, 993–1003 (2006).

11. M. E. Thomason, et al., Weak functional connectivity in the human fetal brain prior to preterm birth. Sci. Rep. 7, 39286 (2017).

12. V. Menon, Developmental pathways to functional brain networks: emerging principles. Trends Cogn. Sci. 17, 627–640 (2013).

13. L. Q. Uddin, K. S. Supekar, S. Ryali, V. Menon, Dynamic reconfiguration of structural and functional connectivity across core neurocognitive brain networks with development. J. Neurosci. Off. J. Soc. Neurosci. 31, 18578–18589 (2011).

14. E. Paraskevopoulos, N. Chalas, P. Bamidis, Functional connectivity of the cortical network supporting statistical learning in musicians and non-musicians: an MEG study. Sci. Rep. 7, 1–10 (2017).

15. B. A. E. Hunt, et al., Relationships between cortical myeloarchitecture and electrophysiological networks. Proc. Natl. Acad. Sci. 113, 13510–13515 (2016).

16. N. Geades, et al., Quantitative analysis of the z-spectrum using a numerically simulated look-up table: Application to the healthy human brain at 7T. Magn. Reson. Med. (2016) https://doi.org/10.1002/mrm.26459.

17. M. F. Glasser, D. C. Van Essen, Mapping Human Cortical Areas In Vivo Based on Myelin Content as Revealed by T1- and T2-Weighted MRI. J. Neurosci. 31, 11597–11616 (2011).

18. M. F. Glasser, et al., The minimal preprocessing pipelines for the Human Connectome Project. Neuroimage 80, 105–124 (2013).

19. M. F. Glasser, M. S. Goyal, T. M. Preuss, M. E. Raichle, D. C. Van Essen, Trends and properties of human cerebral cortex: Correlations with cortical myelin content. Neuroimage 93, 165–175 (2014).

20. D. C. Van Essen, et al., The WU-Minn Human Connectome Project: An overview. Neuroimage 80, 62–79 (2013).

21. M. M. Vandewouw, J. M. Young, M. M. Shroff, M. J. Taylor, J. G. Sled, Altered myelin maturation in four year old children born very preterm. NeuroImage Clin. 21 (2019).

22. Z. Ma, N. Zhang, Cross-population myelination covariance of human cerebral cortex. Hum. Brain Mapp. 38, 4730–4743 (2017).

23. N. Tzourio-Mazoyer, et al., Automated anatomical labeling of activations in SPM using a macroscopic anatomical parcellation of the MNI MRI single-subject brain. Neuroimage 15, 273–289 (2002).

24. L. Melie-Garcia, et al., Networks of myelin covariance. Hum. Brain Mapp. 39, 1532–1554 (2018).

25. F. Váša, et al., Adolescent tuning of association cortex in human structural brain networks. Cereb. Cortex 28, 281–294 (2018).

26. A. M. Fjell, et al., When does brain aging accelerate? Dangers of quadratic fits in cross-sectional studies. Neuroimage 50, 1376–1383 (2010).

27. K. S. Aboud, et al., Structural covariance across the lifespan: Brain development and aging through the lens of inter-network relationships. Hum. Brain Mapp. 40, 125–136 (2019).

28. N. U. F. F. Dosenbach, et al., Prediction of individual brain maturity using fMRI. Science. 329, 1358–1361 (2010).

29. S. Khan, et al., Maturation trajectories of cortical resting-state networks depend on the mediating frequency band. Neuroimage 174, 57–68 (2018).

30. M. J. Brookes, et al., Altered temporal stability in dynamic neural networks underlies connectivity changes in neurodevelopment. Neuroimage 174, 563–575 (2018).

31. B. A. E. Hunt, et al., Spatial and spectral trajectories in typical neurodevelopment from childhood to middle age. Netw. Neurosci. 3, 497–520 (2019).

32. C. B. Schäfer, B. R. Morgan, A. X. Ye, M. J. Taylor, S. M. Doesburg, Oscillations, networks, and their development: MEG connectivity changes with age. Hum. Brain Mapp. 35, 5249–5261 (2014).

33. D. J. Miller, et al., Prolonged myelination in human neocortical evolution. Proc. Natl. Acad. Sci. U. S. A. 109, 16480–16485 (2012).

34. G. L. Baum, et al., Development of structure–function coupling in human brain networks during youth. Proc. Natl. Acad. Sci. U. S. A. 117, 771–778 (2020).

35. C. Paquola, et al., Microstructural and functional gradients are increasingly dissociated in transmodal cortices. PLoS Biol. 17 (2019).

36. B. Vázquez-Rodríguez, et al., Gradients of structure–function tethering across neocortex. Proc. Natl. Acad. Sci. U. S. A. 116, 21219–21227 (2019).

37. A. Von Stein, J. Sarnthein, Different frequencies for different scales of cortical integration: From local gamma to long range alpha/theta synchronization in International Journal of Psychophysiology, (2000), pp. 301–313.

38. M. Siegel, T. H. Donner, A. K. Engel, Spectral fingerprints of large-scale neuronal interactions. Nat. Rev. Neurosci. 13, 121–134 (2012).

39. S. M. Doesburg, J. J. Green, J. J. McDonald, L. M. Ward, Theta modulation of interregional gamma synchronization during auditory attention control. Brain Res. 1431, 77–85 (2012).

40. S. M. Doesburg, S. A. Vinette, M. J. Cheung, E. W. Pang, Theta-modulated gamma-band synchronization among activated regions during a verb generation task. Front. Psychol. 3 (2012).

41. E. Başar, C. Başar-Ero□lu, B. Güntekin, G. G. Yener, “Brain’s alpha, beta, gamma, delta, and theta oscillations in neuropsychiatric diseases: Proposal for biomarker strategies” in Supplements to Clinical Neurophysiology, (2013), pp. 19–54.

42. P. J. Uhlhaas, W. Singer, High-frequency oscillations and the neurobiology of schizophrenia. Dialogues Clin. Neurosci. 15, 301–313 (2013).

43. G. Buzsáki, E. W. Schomburg, What does gamma coherence tell us about inter-regional neural communication? Nat. Neurosci. 18, 484–489 (2015).

44. J. Cannon, et al., Neurosystems: Brain rhythms and cognitive processing. Eur. J. Neurosci. 39, 705–719 (2014).

45. S. Wenzhi, D. Yang, Layer-specific network oscillation and spatiotemporal receptive field in the visual cortex. Proc. Natl. Acad. Sci. U. S. A. 106, 17986–17991 (2009).

46. A. K. Roopun, et al., A beta2-frequency (20-30 Hz) oscillation in nonsynaptic networks of somatosensory cortex. Proc. Natl. Acad. Sci. U. S. A. 103, 15646–15650 (2006).

47. S. Sadaghiani, A. Kleinschmidt, Brain Networks and α-Oscillations: Structural and Functional Foundations of Cognitive Control. Trends Cogn. Sci. 20, 805–817 (2016).

48. O. Jensen, A. Mazaheri, Shaping functional architecture by oscillatory alpha activity: Gating by inhibition. Front. Hum. Neurosci. 4 (2010).

49. O. Jensen, Oscillations in the Alpha Band (9-12 Hz) Increase with Memory Load during Retention in a Short-term Memory Task. Cereb. Cortex 12, 877–882 (2002).

50. A. Dale, B. Fischl, M. I. Sereno, Cortical Surface-Based Analysis: I. Segmentation and Surface Reconstruction. Neuroimage 9, 179–194 (1999).

51. A. M. Dale, M. I. Sereno, Improved localization of cortical activity by combining EEG and MEG with MRI cortical surface reconstruction: A linear approach. J. Cogn. Neurosci. 5, 162–176 (1993).

52. B. Fischl, A. M. Dale, Measuring the thickness of the human cerebral cortex from magnetic resonance images. Proc. Natl. Acad. Sci. U. S. A. 97, 11050–11055 (2000).

53. B. Fischl, A. Liu, A. ∼M. Dale, Automated manifold surgery: constructing geometrically accurate and topologically correct models of the human cerebral cortex. {IEEE} Med. Imaging 20, 70–80 (2001).

54. B. Fischl, M. I. Sereno, A. Dale, Cortical Surface-Based Analysis: II: Inflation, Flattening, and a Surface-Based Coordinate System. Neuroimage 9, 195–207 (1999).

55. B. Fischl, M. I. Sereno, R. B. H. Tootell, A. M. Dale, High-resolution intersubject averaging and a coordinate system for the cortical surface. Hum. Brain Mapp. 8, 272–284 (1999).

56. B. Fischl, et al., Whole brain segmentation: automated labeling of neuroanatomical structures in the human brain. Neuron 33, 341–355 (2002).

57. B. Fischl, et al., Sequence-independent segmentation of magnetic resonance images. Neuroimage 23, S69–S84 (2004).

58. X. Han, et al., Reliability of MRI-derived measurements of human cerebral cortical thickness: the effects of field strength, scanner upgrade and manufacturer. Neuroimage 32, 180–194 (2006).

59. J. Jovicich, et al., Reliability in multi-site structural MRI studies: Effects of gradient nonlinearity correction on phantom and human data. Neuroimage 30, 436–443 (2006).

60. M. Reuter, H. D. Rosas, B. Fischl, Highly Accurate Inverse Consistent Registration: A Robust Approach. Neuroimage 53, 1181–1196 (2010).

61. M. Reuter, N. J. Schmansky, H. D. Rosas, B. Fischl, Within-Subject Template Estimation for Unbiased Longitudinal Image Analysis. Neuroimage 61, 1402–1418 (2012).

62. M. Ganzetti, N. Wenderoth, D. Mantini, Whole brain myelin mapping using T1- and T2- weighted MR imaging data. Front. Hum. Neurosci. 8, 671 (2014).

63. M. Jenkinson, S. Smith, A global optimisation method for robust affine registration of brain images. Med. Image Anal. 5, 143–156 (2001).

64. M. Jenkinson, P. Bannister, M. Brady, S. Smith, Improved optimisation for the robust and accurate linear registration and motion correction of brain images. Neuroimage 17, 825–841 (2002).

65. J. G. Sled, A. P. Zijdenbos, A. C. Evans, A nonparametric method for automatic correction of intensity nonuniformity in MRI data. IEEE Trans. Med. Imaging 17, 87–97 (1998).

66. J. L. R. Andersson, M. Jenkinson, S. Smith, “Non-linear registration, aka spatial normalisation. FMRIB Technial Report TR07JA2.” (2007).

67. M. W. Woolrich, et al., Bayesian analysis of neuroimaging data in FSL. Neuroimage 45, S173–S186 (2009).

68. S. D. Muthukumaraswamy, High-frequency brain activity and muscle artifacts in MEG/EEG: a review and recommendations. Front. Hum. Neurosci. 7 (2013).

69. B. B. D. Van Veen, W. Van Drongelen, M. Yuchtman, A. Suzuki, Localization of brain electrical activity via linearly constrained minimum variance spatial filtering. IEEE Trans. Biomed. Eng. 44, 867–880 (1997).

70. R. Oostenveld, P. Fries, E. Maris, J. M. Schoffelen, FieldTrip: Open source software for advanced analysis of MEG, EEG, and invasive electrophysiological data. Comput. Intell. Neurosci. 2011, 1 (2011).

71. J. Sarvas, Basic mathematical and electromagnetic concepts of the biomagnetic inverse problem. Phys. Med. Biol. 32, 11 (1987).

72. G. Nolte, The magnetic lead field theorem in the quasi-static approximation and its use for magnetoencephalography forward calculation in realistic volume conductors. Phys. Med. Biol. 48, 3637 (2003).

73. A. N. Tikhonov, On the stability of inverse problems. Comptes Rendus L Acad. Des Sci. L Urss 39, 176–179 (1943).

74. G. L. Colclough, M. J. Brookes, S. M. Smith, M. W. Woolrich, A symmetric multivariate leakage correction for MEG connectomes. Neuroimage 117, 439–448 (2015).

75. R. Bartoszy□ski, M. Niewiadomska-Bugaj, Probability and Statistical Inference (2007) https://doi.org/10.1002/9780470191590.

76. P. T. Reiss, et al., Massively parallel nonparametric regression, with an application to developmental brain mapping. J. Comput. Graph. Stat. 23, 232–248 (2014).

77. Y. Benjamini, Y. Hochberg, Controlling the false discovery rate: A practical and powerful approach to multiple testing. J. R. Stat. Soc. 57, 289–300 (1995).

