## Supplemental Information for "The developing relations between networks of cortical myelin and neurophysiological connectivity"

Marlee M. Vandewouw*^^1,2^, Benjamin A.E. Hunt^^1,2^, Justine Ziolkowski^2^, Margot J. Taylor^1,2,3,4^

^1^ Department of Diagnostic Imaging, Hospital for Sick Children, Toronto, Canada

^2^ Program in Neurosciences and Mental Health, Hospital for Sick Children, Toronto

^3^ Department of Psychology, University of Toronto, Toronto, Canada

^4^ Department of Medical Imaging, University of Toronto, Toronto, Canada

^These co-authors contributed equally to the work.

* Corresponding author:

Marlee M. Vandewouw

Diagnostic Imaging, Hospital for Sick Children,

555 University Ave., Toronto, ON M5G 1X8, Canada

**Sliding window parameter variation**

The following supplemental figures demonstrate the invariance of our results to the sliding window parameters. Results in the main text are presented with a sliding window width (*w*) of 50 and a step size (*s*) of 2. In each of the following figures, the results presented in Figures 1-4 in the main text are replicated with step sizes of 1 and 5 (Supplemental Figures 1, 3, 5, and 7), and window widths of 40 and 60 (Supplemental Figures 2, 4, 6, and 8).

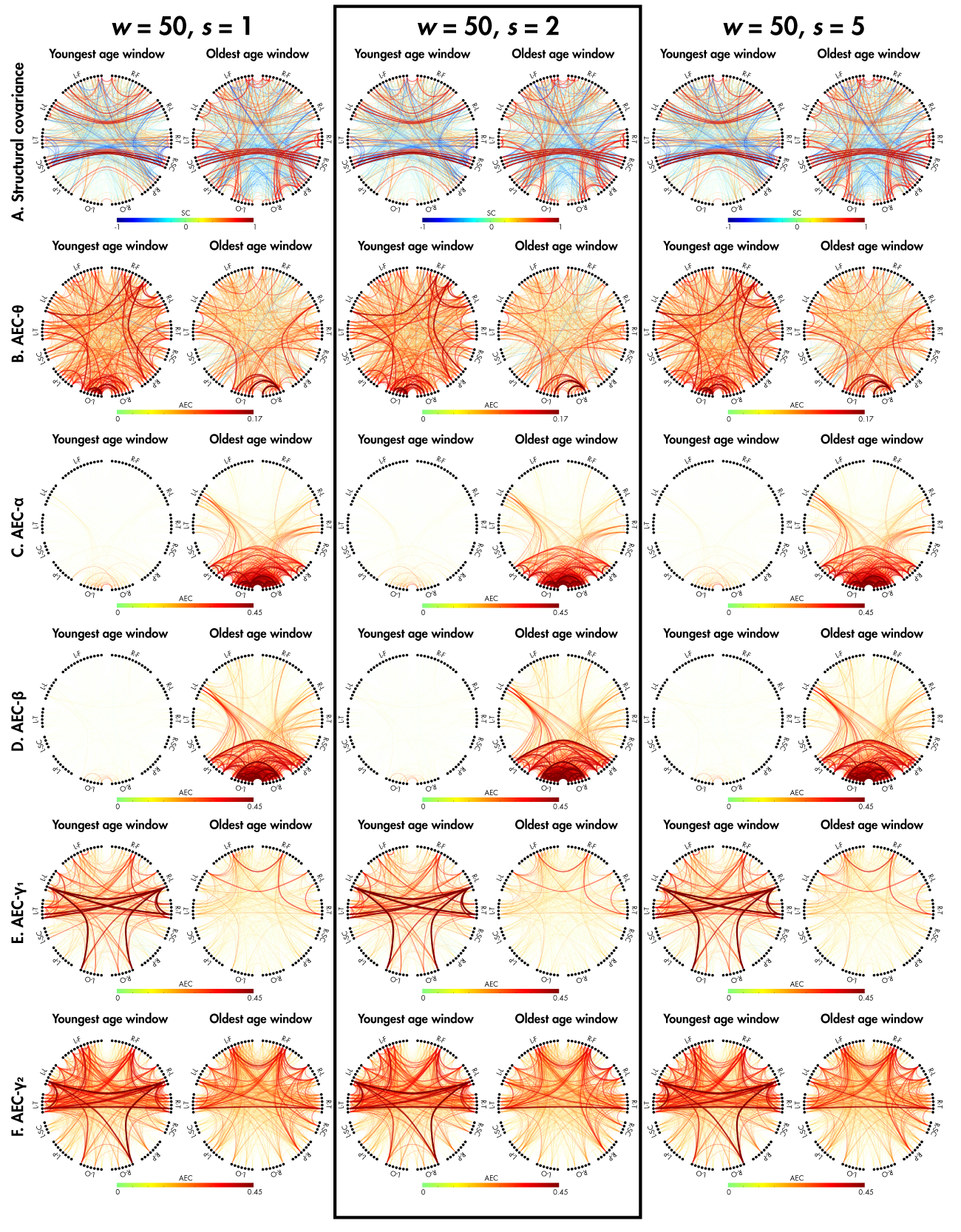

**Supplemental Figure 1.** Replication of Figure 1 from the main text, with step size varied between 1, 2, and 5. Results corresponding to those presented in the main text are shown in the black box.

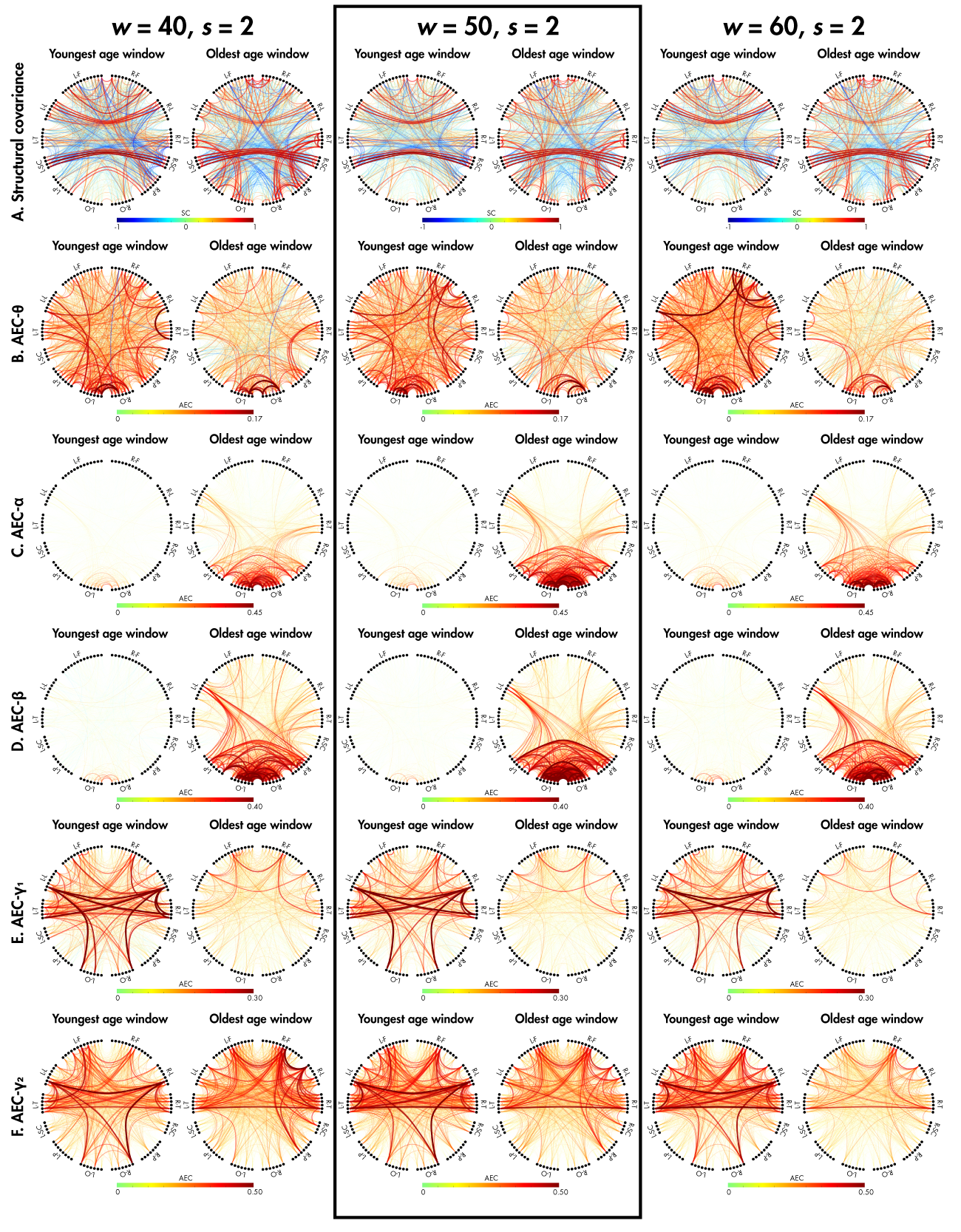

**Supplemental Figure 2.** Replication of Figure 1 from the main text, with window width varied between 40, 50, and 60. Results corresponding to those presented in the main text are shown in the black box.

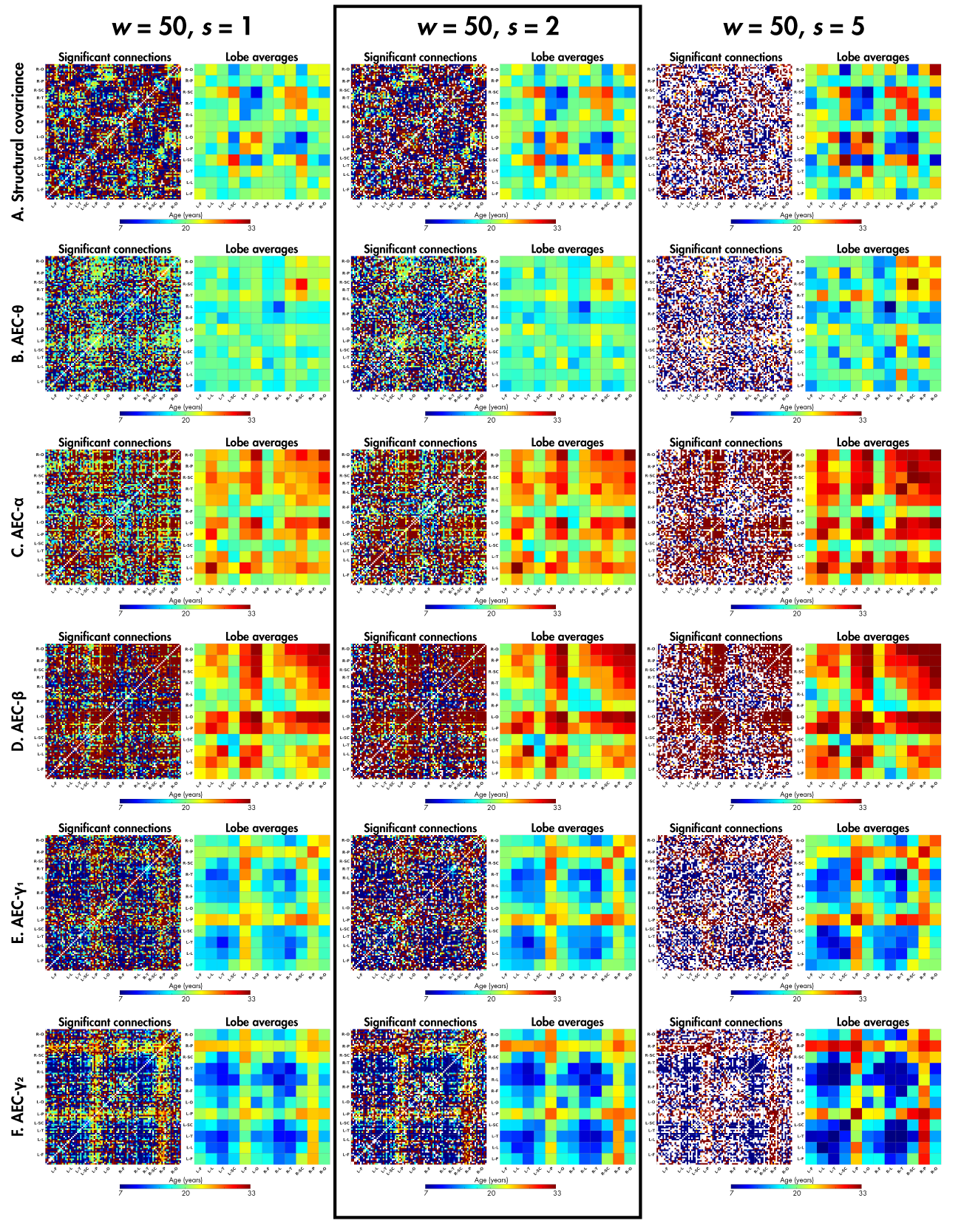

**Supplemental Figure 3.** Replication of Figure 2 from the main text, with step size varied between 1, 2, and 5. Results corresponding to those presented in the main text are shown in the black box.

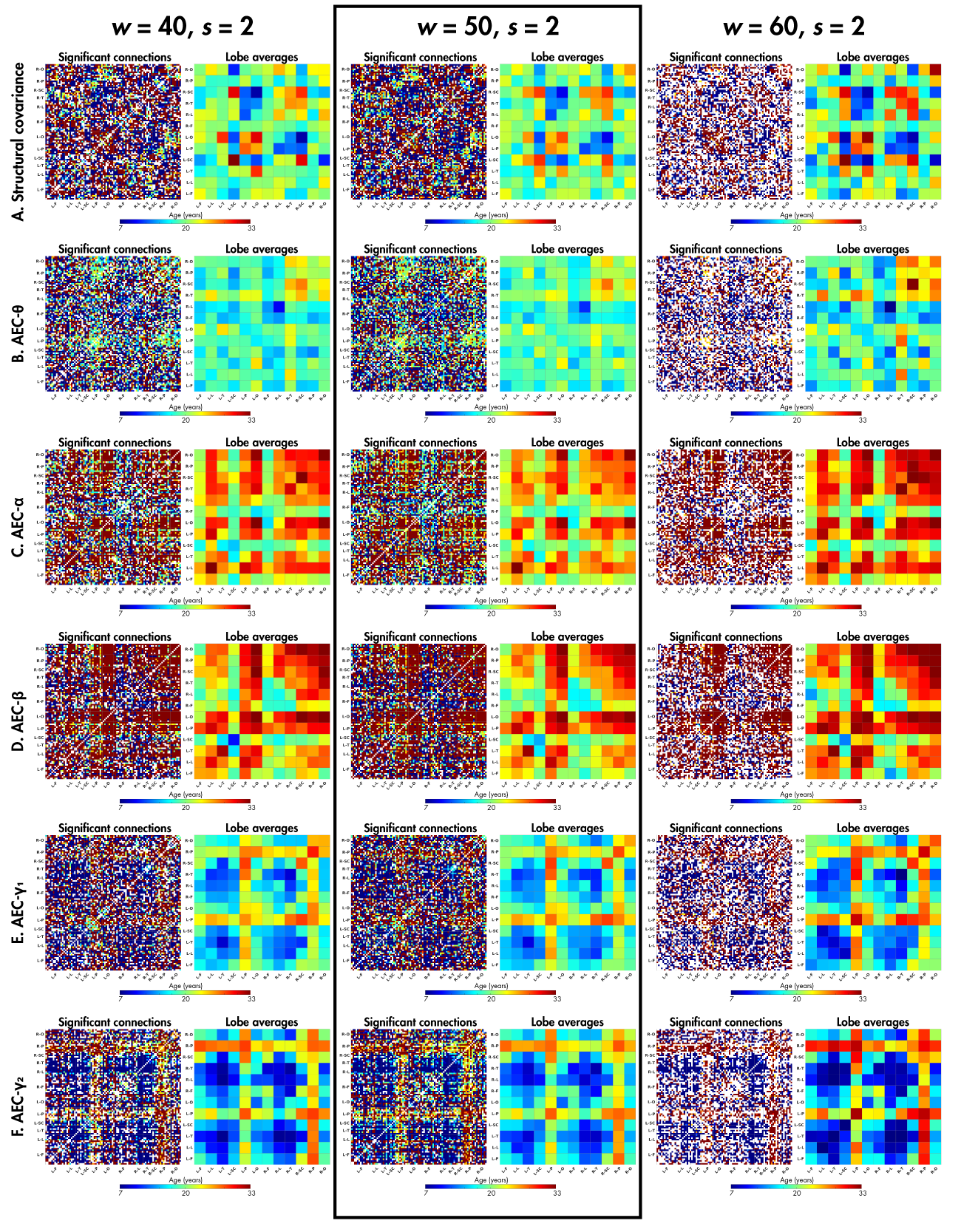

**Supplemental Figure 4.** Replication of Figure 2 from the main text, with window width varied between 40, 50, and 60. Results corresponding to those presented in the main text are shown in the black box.

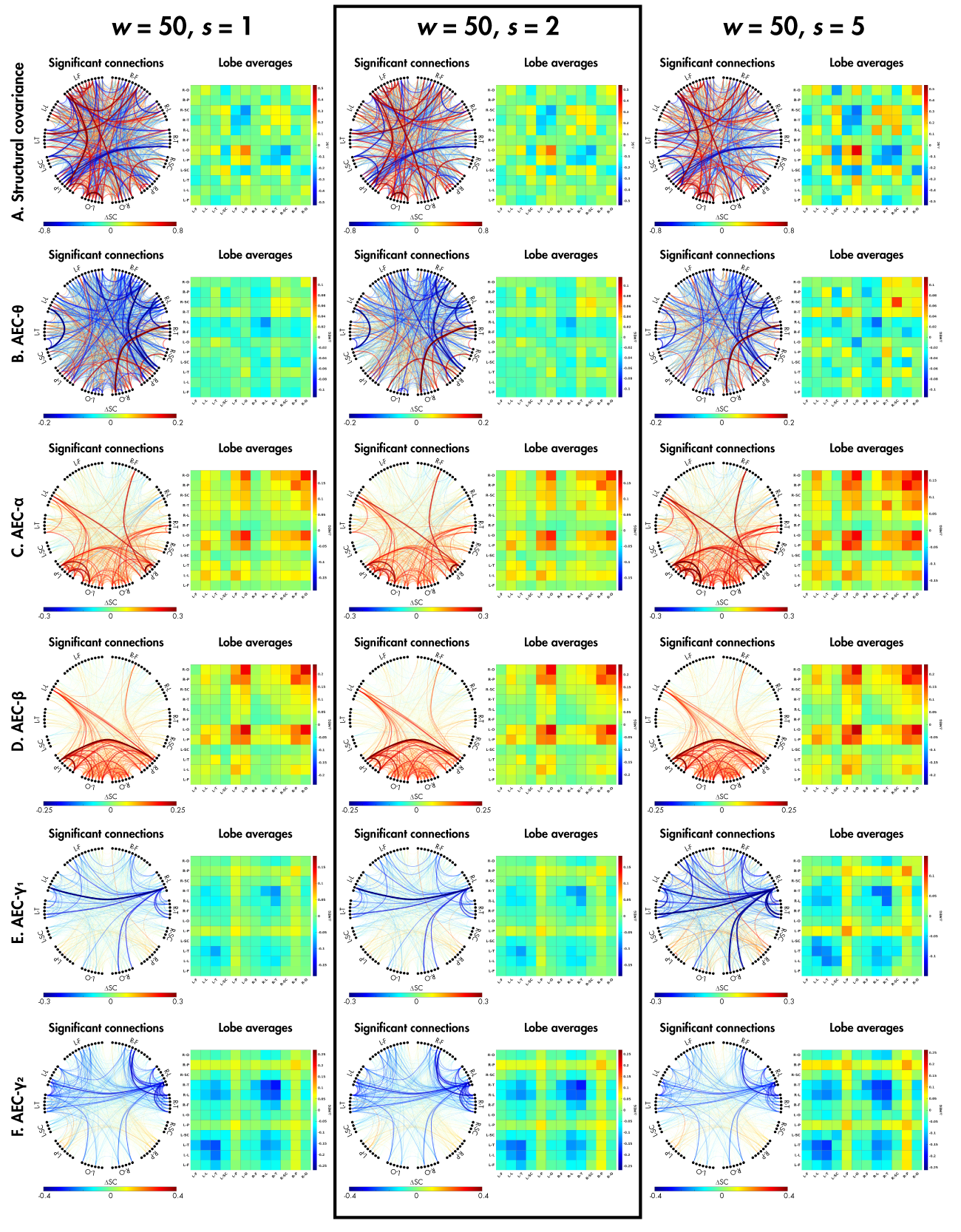

**Supplemental Figure 5.** Replication of Figure 3 from the main text, with step size varied between 1, 2, and 5. Results corresponding to those presented in the main text are shown in the black box.

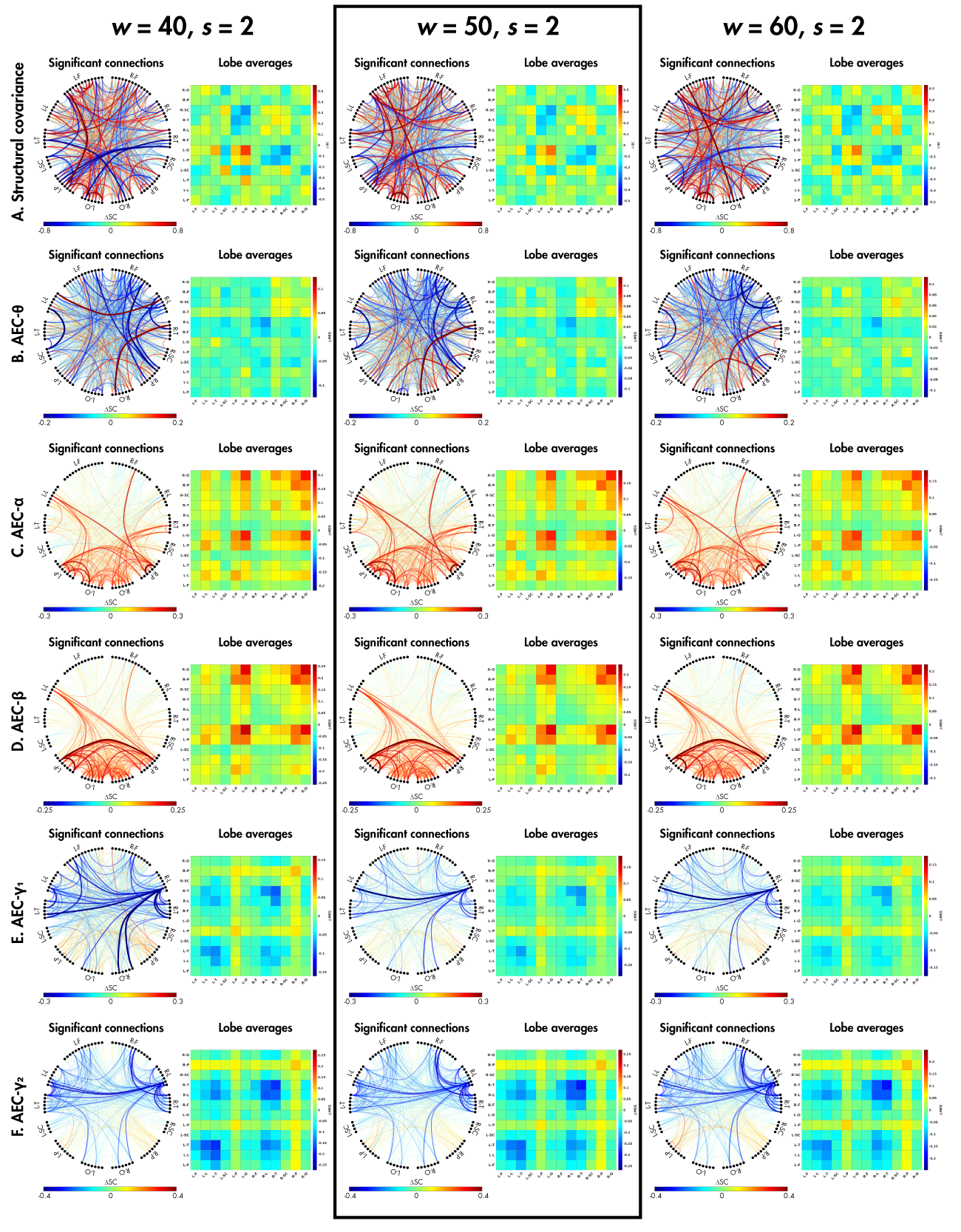

**Supplemental Figure 6.** Replication of Figure 3 from the main text, with window width varied between 40, 50, and 60. Results corresponding to those presented in the main text are shown in the black box.

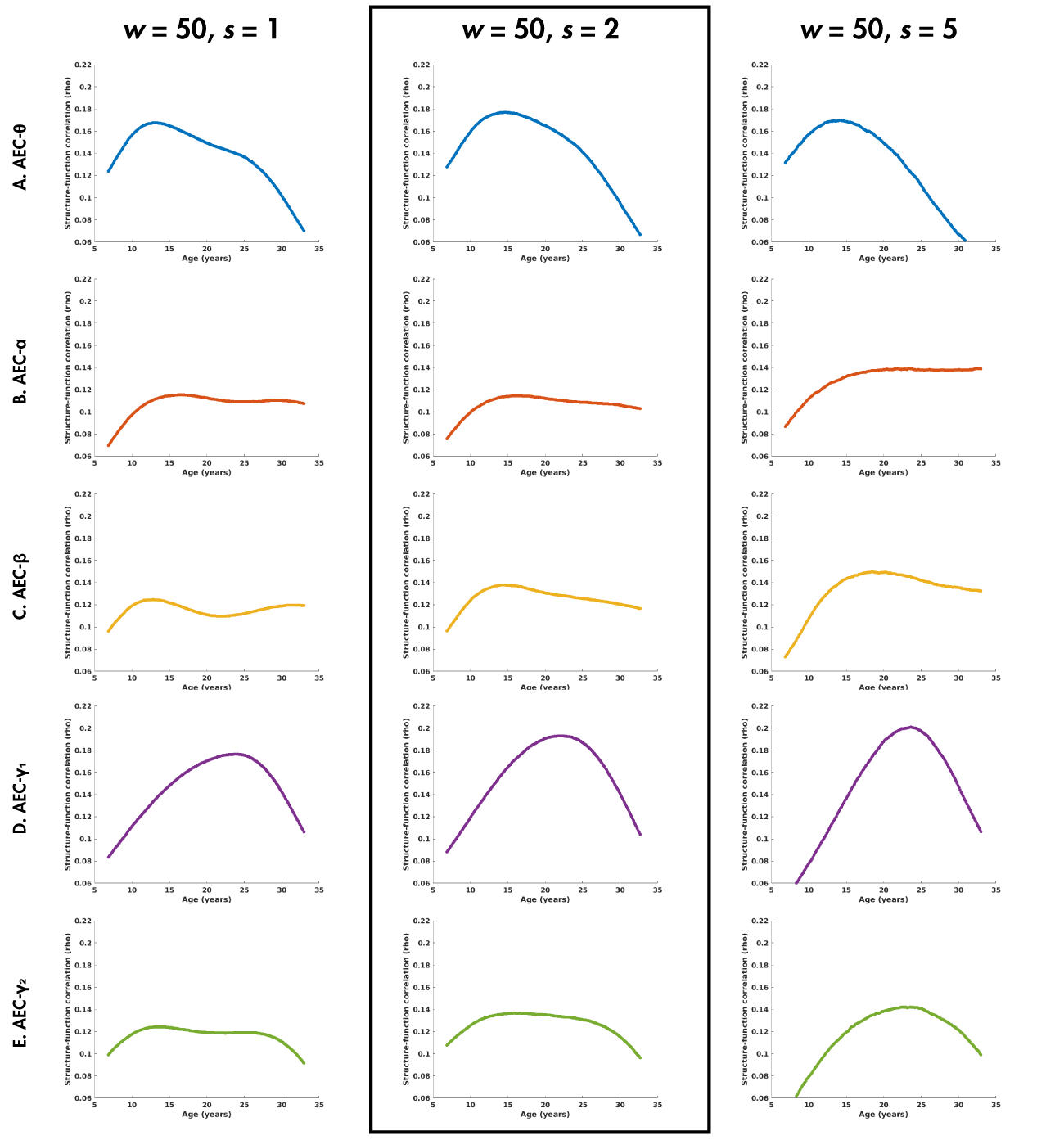

**Supplemental Figure 7.** Replication of Figure 4 from the main text, with step size varied between 1, 2, and 5. Results corresponding to those presented in the main text are shown in the black box.

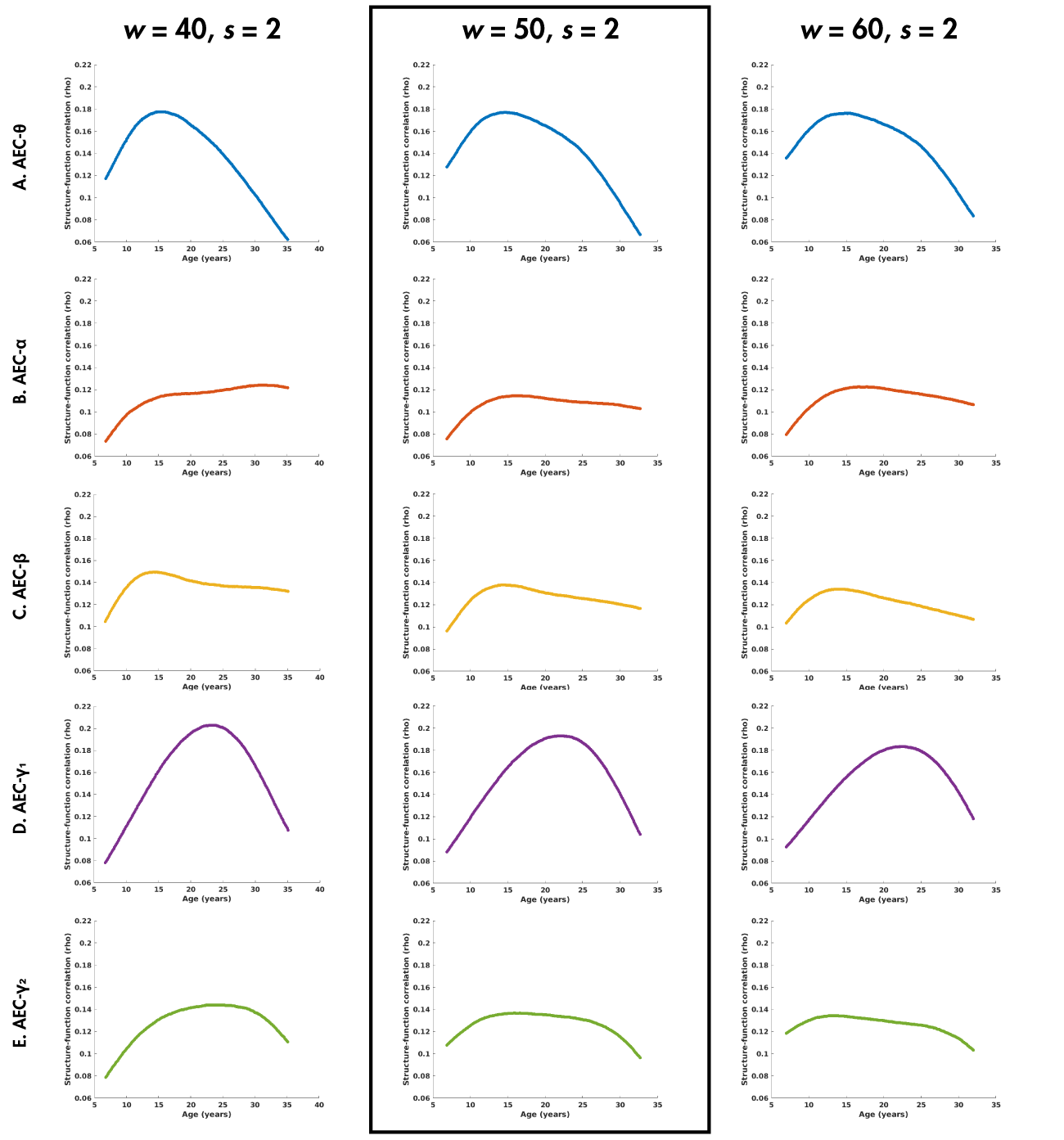

**Supplemental Figure 8.** Replication of Figure 4 from the main text, with window width varied between 40, 50, and 60. Results corresponding to those presented in the main text are shown in the black box.
